# Multi-faceted role of dopamine afferents to the ventral pallidum in reward processing

**DOI:** 10.64898/2026.11.03.763967

**Authors:** Matthew William Spooner, Jennifer L. Cornish, Asheeta A. Prasad

**Affiliations:** Discipline of Neuroscience, School of Medical Sciences, Faculty of Medicine and Health, University of Sydney, Sydney, NSW, Australia; School of Psychological Sciences, Faculty of Medicine, Health and Human Sciences, Macquarie University, Sydney, NSW, Australia

**Keywords:** ventral pallidum, dopamine, reward, learning, motivation

## Abstract

The ventral pallidum (VP) is necessary for reward processing and a key node in the brain’s dopamine (DA) pathways. Yet the direct DA inputs have not been studied during reward learning. Here, we elucidate the role of VP DA in reward processing using circuit tracing, fibre photometry, and behavioural pharmacology in male rats. We show that VP receives both non-DA and DA inputs from the midbrain. During an instrumental reward conditioning paradigm, we characterized VP DA activity using a DA sensor (GRAB_DA1h) across learning, extinction, re-acquisition, and motivational test. Followed by pharmacological manipulation using the DA-antagonist (fluphenazine) in VP to demonstrate a causal role in reward learning and motivation. Together our findings advance the functional contributions of non-canonical mesolimbic DA pathways.

## Introduction

Dopamine (DA) is the primary neurotransmitter involved in reward learning, reward expectation, and motivation^1,2^. In appetitive instrumental conditioning, DA supports the acquisition of action-outcome associations^3,4^, motivation to work for rewards ^5^ and drives learning-related plasticity (Shen et al., 2008). These multi-faceted roles of dopamine are dependent on dopamine neuronal activity and dopamine release in the various projection regions^6^. Previous studies show dopamine sends afferents to the ventral pallidum (VP) ^7,8^. Moreover, expression of dopamine receptors in the VP has been evolutionarily conserved across species^9^. In rodents, electrophysiological studies have demonstrated that both dopamine receptors D1 and D2 are present in the VP and can independently activate neuronal activity^7,10^.

The functions of VP are centred around hedonic behavior^9,11^. For example, lesions or pharmacological inhibition of the VP reduce voluntary food consumption and hedonic responses to food rewards^12^, curb willingness to work for sucrose rewards ^13^ and impair instrumental conditioning for drugs like cocaine ^14^ and alcohol ^15,16^. Along with others, we have previously shown the importance of VP and specific VP cell types, and VP afferents in relapse behaviour^16–20^. Findings from several studies have emphasized the important functions of DA receptors, including VP DRD3 efferents^21–24^. However, the direct role of DA in the VP remains poorly elucidated. To advance our understanding of DA functions in VP, we examined two overlooked aspects of VP reward processing; firstly, the different sources of dopaminergic input into the VP, and second, the role DA inputs play in the VP across different stages of reward processing behaviour (reward acquisition, extinction, and re-acquisition).

DA likely supports reward learning in the VP, which is innervated by midbrain DA cells^7,8^. VP neurons also exhibit similar functional properties to DA activity, with neural activity consistent with reward value and prediction errors^25–28^. Although several studies have reported the functional connectivity between midbrain and VP, the role of DA inputs to the VP in the reward processing has not been directly examined^16,29,30^. To address this gap, we first confirmed the neural DA afferents to the VP. Next, we performed fibre photometry to record DA activity in the VP during an appetitive instrumental conditioning task. We found DA signals in the VP tracked reward acquisition, extinction, reacquisition, and motivated behaviours. Critically, pharmacological blockade of these signals impaired reward learning and reduced motivation, demonstrating their causal role in driving reward learning and motivation. As such, we reveal that VP DA supports reward processing and instrumental reward learning and drives motivation to work for rewards.

## Materials and Methods

### Subjects

This study used male Sprague-Dawley rats (250-350g, 7 weeks old on arrival), obtained from the Animal Resource Centre (Perth, Australia). N=4 for tracing study, n=10 for fibre photometry experiments and n=25 for pharmacology experiments (n=15 for self-administration and n=10 for locomotor). Rats were housed in a climate-controlled colony room with a 12hr light-dark cycle and 2-4 animals per cage. All behavioural procedures were conducted in the light cycle. Subjects had *ad libitum* access to food until the experiment began, after which each rat received 20g of food per day. Procedures were approved by the University of Sydney’s Animal Ethics Committee. They adhered to the National Health and Medical Research Code for the Care and Use of Animals for Scientific Purposes in Australia (2013).

### Retrograde tracers and viral vectors

For circuit mapping, the retrograde tracer cholera toxin b subunit (CTb) conjugated to AlexaFluor-488 (CTb-488, #C34775 ThermoFisher Scientific) was used to target VP afferents. For dual accumbens and VP circuit tracing, AlexaFluor-488 (CTb-488) was placed in the VP and AlexaFluor-555 (CTb-555) (#C34776, ThermoFisher Scientific) in accumbens. CTB tracers were diluted to 1.0 mg/mL and 40nl was injected per brain region. For fibre photometry, viral vectors were AAV-hsyn-GRAB_DA1h (750nl, Addgene viral prep #113050-AAV9; http://n2t.net/addgene:113050; RRID:Addgene_113050, titer ≥ 1×10¹³ vg/mL) for the dopamine sensor.

### Behavioral procedures

#### Magazine training

During the first 2 days of training, animals received 2 sessions of magazine training each day, with 10 pellets per session dispensed at random intervals. Each session was 20 min, and sessions in all subsequent stages were 30 min. *Lever press training:* For the next 10 days, rats were trained to discriminate between an active lever that provided sugar pellet reward with an FR1 schedule and with a 10 s timeout period and an inactive lever that led no rewards. *Extinction:* For the following 4 days, rats underwent extinction, during which neither lever provided any sugar pellet rewards.

#### Reacquisition

Rats then underwent 1 day of retraining, during which the active lever once again gave sugar pellets at an FR1 schedule with followed by a 10 s timeout. If there was a drug effect during pharmacological experiments, rats were trained until drug, and control groups were matched for performance. *Progressive ratio test:* In the final stage, all rats underwent a progressive ratio (PR) test of their motivation to lever press for sugar pellets ^16^. During this test, similar all sessions, both the active and inactive lever are always present and but rewarded on a PR schedule of active lever presses. The schedule was derived from the function where response ratio equals to [5e(reinforcer number × 0.2)] − 5, rounded to the nearest integer ^31^. During this test, the ratio of lever presses to pellet delivery progressively increased, thereby assessing their willingness to work for rewards.

### Training apparatus

Standard rat operant chambers (Med Associates) with dimensions 29.5 cm (width) × 24.8 cm (length) × 18.7 cm (height) were used for all training procedures. Chambers were also located inside wooden cabinets to attenuate external light and sound. The chambers contained two levers, which are always extended. each symmetrically located on one wall of the chamber, 3 cm above the grid floor. A recessed magazine was located behind a 4 × 4 cm opening in the centre of the same wall, positioned between the two levers.

Pressing the active lever delivered a sugar pellet (45mg, Able Scientific, product code: ASF06162) to the magazine during instrumental conditioning, with an intertrial interval (ITI) of 10s. Pellets were delivered on a fixed-ratio (FR1) schedule. By contrast, responding to the inactive lever triggered no events. A computer running MedPC-IV software controlled all events.

### Surgeries and infusions

Rats were anesthetized with isoflurane (5%) and placed in a stereotax (Kopf Instruments). Anaesthesia was maintained at 1.5-3% until the completion of surgery. Subcutaneous injections of carprofen (5mg/kg, Wagg & Purr), Duplocillin (0.15mL/kg, MSD Animal Health), and cefazolin (75mg/kg, AFT Pharmaceuticals) were given to relieve pain and prevent infection. Bupivacaine (0.1mL, 0.5%, Pfizer) was administered at the incision site for local anaesthesia. Injections of tracers and viral vectors targeting the VP used coordinates: A/P = 0.00; M/L = ± 2.3; D/V = −8.2 and accumbens A/P = 1.92; M/L = ± 1.1; D/V = −6.2 ^32^. Guide cannulas (11mm, RWD) were implanted at a 10° angle for cannulation experiments. Using trigonometry, the change in angle yielded the following coordinates: A/P = 0.2; M/L = ±3.75; D/V = −7.8. Surgical procedures for cannula implantations are similar to described in Xie and Prasad 2022 ^33^. Rats infused with tracers were given 1 week to recover and a further 3 weeks for the tracers to retrogradely disperse. For fiber photometry, behavioral procedures began 6 weeks post-surgery to allow for recovery and viral expression. For cannulation studies involving pharmacological interventions, rats were given a minimum of 1 week to recover before behavioral procedures began.

### Fiber photometry Recordings

Fiber photometry recordings were conducted on instrumental conditioning days 3 (early learning) and 10 (late learning), extinction days 1 (early extinction) and 3 (late extinction), reacquisition, and the progressive ratio (i.e., motivation) test. These recording days were chosen to compare early and established phases of reward learning and extinction, reacquisition, and motivation under increasing work requirements. Prior to recording session, animals were habituated to tethering for one to two days to reduce impact of tethering on behaviour. Other studies under these conditions have demonstrated robust acquisition, extinction, and reacquisition of behaviour^16,34–36^. Subjects were tethered to the setup via a cable (Ø400µm, 0.57NA, Doric Lenses) that was fitted to their cannula using a mating sleeve (ADAF1-5, ThorLabs). A real-time signal processor (RZ10X Lux-I/0 Processor, Tucker-Davis Technologies) transmitted light at 465nm and 405nm to excite GRAB-DA1h. The proteins have two conformations: bound and unbound (Sun et al., 2020; Zhang et al., 2023). The 465nm LED stimulated dopamine-binding dependent fluorescent signals, while the 405nm LED was used for dopamine-independent fluorescence. Consistent with standard protocol, the experimental control was the 405nm signal, that provided a negative internal isosbestic control^37^.

The LED driver sinusoidally modulated LEDs at 209Hz for the 465nm LED and 331Hz for the 405nm LED, maintaining light intensity between 10-30 µW to minimize photo-bleaching. Driver cables (Ø200µm, 0.5 NA, LXFX-KIT-2C, Tucker-Davis Technologies) sent light through the Doric Mini Cube (FMC6_IE, Doric Lenses) to the subject. Fluorescence was collected and transmitted to the RZ10x photosensor via response cables (Ø600µm, 0.5 NA, LXFX-KIT-2C, Tucker-Davis Technologies). The photosensor low-pass filtered the data at 6Hz to remove noise and demodulated 405nm and 465nm signals. Synapse software (Tucker-Davis Technologies) controlled the RZ10x and recorded signals. Behavioral events were time-locked to fiber photometry data by Synapse.

### Fibre photometry data analysis

Custom MATLAB (Mathworks) scripts were employed for data analysis. Time ranges in the raw data stream, with motion artifacts, were manually excluded. High-frequency noise was removed through low-pass digital filtering at 2 Hz. To control for dopamine-independent fluorescence and to normalize data, ΔF/F (change in fluorescence relative to baseline) was calculated^37^. The control 405nm signal was linearly fit to the 465nm signal. ΔF/F was calculated with the formula (465nm signal −fitted 405nm signal) / (fitted 405nm signal), yielding the final signal.

For each subject and recording session, dopamine signals were Z-scored around behavioral events: LP+ (lever press resulting in reward delivery) and LP− (lever press with no reward delivery during 10-second time-out). Events were isolated temporally such that photometry signal did not overlap between examined events. Z-scores were calculated on a trial-by-trial basis using the formula (all data points ΔF/F - baseline mean ΔF/F) / (baseline standard deviation ΔF/F) where baseline was 10 to 5-seconds pre-behavior and all data points were ± 10-seconds peri-behavior. Group mean of the Z-score signal surrounding behavioral events was determined.

The area-under-the-curve (AUC) of group mean Z-scores was computed with trapezoidal numerical integration to examine behavior-related increases in activity or dopamine release. AUCs were calculated in −5 to −3 seconds before a behavior to indicate a ‘baseline’ value, and 0 to 2 seconds after behavioral onset to indicate a ‘response’ value. Data was plotted with MATLAB.

### Pharmacological inhibition of dopamine receptors in the VP

Fluphenazine dihydrochloride (product number: F0280000; molecular weight = 510.44) was purchased from Sigma Aldrich (Castle Hill, NSW, Australia). Rats were allocated to drug or control groups, matched for performance prior to infusion days. All rats receive the same treatment on all days. Fluphenazine was dissolved in 0.9% saline, and the vehicle was 0.9% saline. Fluphenazine was administered at a concentration of 10nmol/500nl, and 500nl per hemisphere was infused at 250nl/min. Infusion needles (33 gauge, 11.5mm, RWD) were connected via polyethylene tubing (RWD) to a 5µl Hamilton syringe, and microinfusions were driven by a pump (Harvard Apparatus, USA). Dose of fluphenazine was based on previous publication^38^. After the two-minute infusion, infusion needles were left in place for a further two minutes to allow for diffusion. Control rats received infusions of 0.9% saline with the same volume, rate, and diffusion time.

### Behavioral data analysis

Across training and testing, data were analyzed using a contrast testing procedure. For the photometry experiment, in which there was a single group, data were analyzed with a within-subjects factor of group and within-subjects factor of trial. For the pharmacological experiment with two groups, data were analyzed using a between-subjects factor for group and a within-subjects factor for trial. The criterion for rejection of the null hypothesis (α) was set at 0.05. For all significant between-group differences, we calculated standardized 95% confidence intervals (95% CI) as well as Cohen’s d (d) as a measure of effect size. By convention, a Cohen’s d of 0.2 indicates a small effect, 0.5 indicates a medium-sized effect, and 0.8 or above indicates a large effect.

### Locomotor analysis

Locomotor activity was assessed in a Noldus Phenotyper (length = 43.2 cm, width = 43.2 cm, height = 30.5 cm). The movement tracked multiple motor behaviours during the session, including ambulatory time, counts, distance moved, resting and vertical time. Ethovision XT software (Noldus) recorded total distances and average speeds. Animals were administered with vehicle and fluphenazine across two day in a counterbalanced manner. Within group analysis was conducted using Paired Sample T-Test.

### Retrograde tracer cell counts and analysis

CTb-488 was infused into the VP to delineate midbrain afferents and stained for tyrosine hydroxylase (TH). Co-labelled TH and CTB quantification revealed dopamine inputs to the VP. Midbrain dopamine neurons were immunostained for TH and the presence of neural tracer, CTb-488. Cells that were positive for either or both of these markers were then counted, and the quantity of cells positive for both TH and CTb-488 indicated connectivity between the VP and midbrain.

### Perfusions and histology

A minimum of 24-hours after behavioral testing completion, subjects were anesthetized with Lethabarb (100mg/kg, i.p., Virdac) and perfused transcardially with 150ml of 0.9% saline then 500ml of 4% paraformaldehyde in 0.1M phosphate buffer, pH 7.4. Brains were removed and post-fixed in 4% paraformaldehyde in 0.1M phosphate buffer for 24 hours at 4°C. The brains were transferred to 30% sucrose in 0.1 M phosphate-buffered saline (PBS) and stored at 4°C for 3 days. Brains were sectioned coronally at 40μm with a cryostat (NX50, Cryostar Expredia). Sections were incubated overnight with a primary anti-TH antibody for labelling DA neurons (mouse anti-tyrosine hydroxylase, 1:500; Cat. AB152, Sigma-Aldrich) or GFP polyclonal antibody for AAV expression primary (chicken anti-GFP antibody, 1:500; GFP polyclonal antibody, AB13970, Invitrogen). Primary antibody was incubated over night at 4 and washed with TBS with 0.05% Tween before incubation with secondary antibody (1:500, Alexa Fluor 488 goat anti-chicken antibody, A-11039, or donkey anti-mouse Alexa Fluor 647, 1:500, Invitrogen Cat. A31571) for 3-hours at room temperature. Sections were counterstained with Hoechst (1:1000, Hoechst 33342, 62249, ThermoFisher Scientific) or DAPI to visualize cell nuclei. Sections were mounted onto Superfrost glass slides (EPBRSF41296, Expredia) and cover-slipped with 100µl mounting media (Prolong Diamond Antifade Mountant, P36970, Invitrogen).

Images were captured with Leica THUNDER (Leica Microsystems) at 20x magnification. For DA efferents to VP analysis, images were analysed using QuPath-0.6.0-rc3-x64 software. Counts of all neurons IR for TH and CTb-488 fluorescence was quantified across three VP sections per animal, n=4. All quantifications were conducted under blind conditions. Delineation of the ventral midbrain area was guided by TH-positive cells and surrounding anatomical landmarks, such as the red nucleus and parabrachial pigmented nucleus. The counting criteria were as follows, TH+ and CTB+ cells must be co-stained with DAPI.

Viral expression and cannula locations were validated using the VP anatomical localisation described by Paxinos and Watson (2006). For experiments using implanted metal cannulae for behavioral pharmacology, once behavioral testing was complete, rats were euthanized, and brains were then sliced on a cryostat to 40-µm thick coronal slices. Every second slice was thawed, mounted on a glass microscope slide, and stained with cresyl violet. Slides were observed under a microscope to confirm the location of cannula tips, and rats with inaccurate cannula placements (one or both cannulas positioned outside the boundaries of the VP) were excluded from the statistical analysis.

## Results

### DA afferents in VP

To identify dopamine afferents to the VP, a neural tracer CTB-488 was placed in the VP (*n*=4), followed by immunohistochemical labelling of dopamine neurons using tyrosine hydroxylase (TH) antibody. Dual quantification of CTB-labelled cells and TH-positive cells in the midbrain was performed to determine the number of DA cells projecting to the VP. We found that 36.02 ± 18.69% of CTB-labelled cells were also TH-positive, while 63.97 ± 18.69% were not TH-positive. Demonstrating both DA (∼36%) and non-DA (∼64%) projections from the midbrain to the VP (Figure 1A-C), similar to previous reports of ∼40% of VP projecting midbrain neurons are TH-positive ^8,39^. As the connectivity between DA neurons and VP has been previously reported, our tracing studies guide the cannula placements for DA recordings and manipulations in the VP.

**Figure 1.**
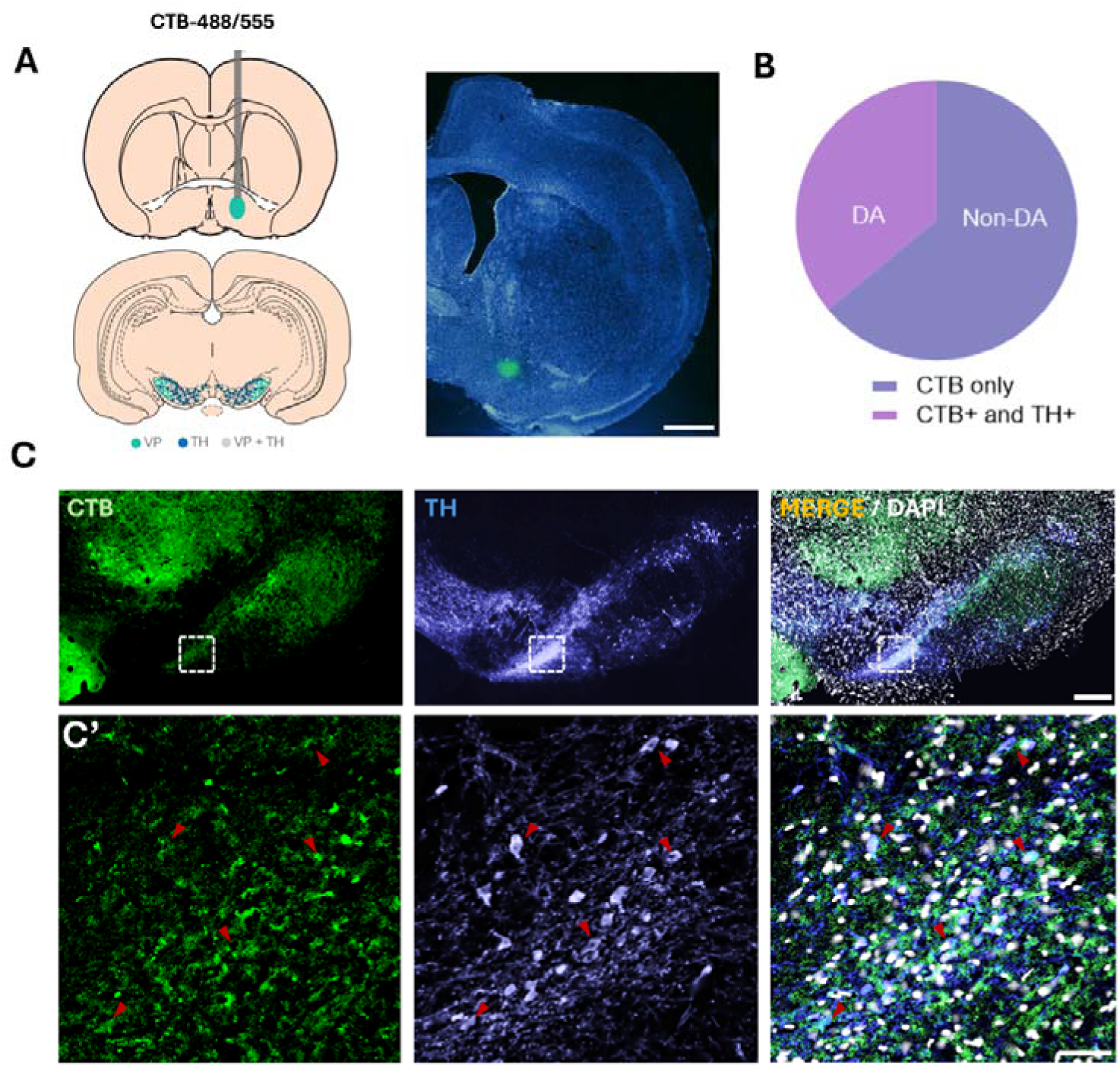
(A) Diagram of surgical procedure to bilaterally infuse CTb into the VP. (B) Representative image showing labelling of VP-projecting midbrain DA neurons. (C) Descriptive statistics showing the percentage of midbrain DA neurons that project to the VP, n=4.

### Histology

Expression of GRAB-DA was determined by immunohistochemistry of GFP fluorescence, and rats were included if the tip of the optic cannula was in the VP *(n* = 10, with one rat excluded due to cannula misplacement). Viral expression and cannula locations were validated using the VP anatomical localisation described by Paxinos and Watson (2006). Placement of optic cannula tips are indicated in Figure 2A-B.

**Figure 2.**
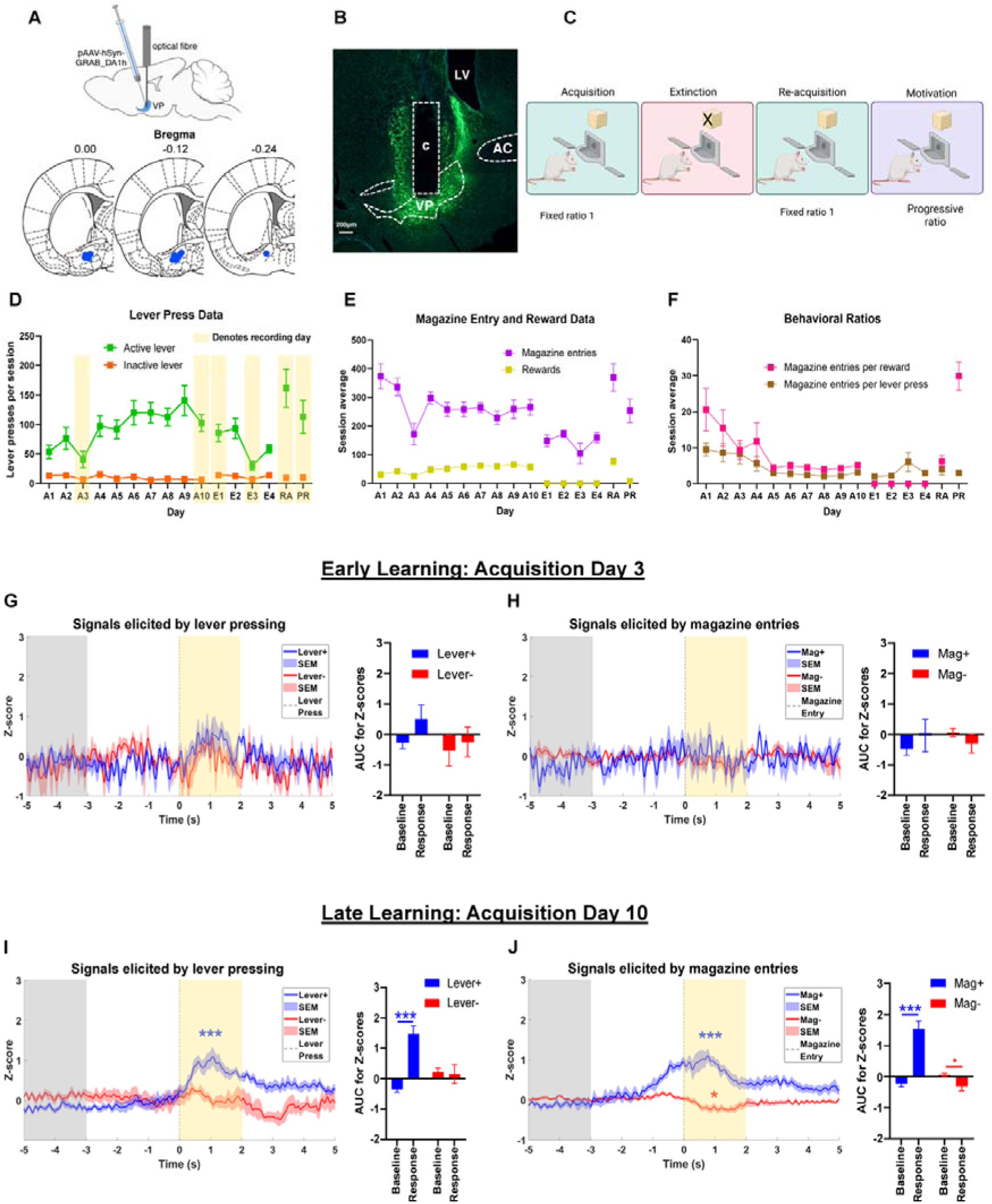
(A) Diagram of surgical procedure of unilateral infusion of GRAB_DA1h and fibre optic cannula implanted into the VP, and histological reconstruction of cannula placement. (B) Representative image of viral spread and cannula placement in the VP, n=10. (C) Diagram of behavioral protocol. (D-F) Mean (± SEM) behavioral data for (D) active and inactive lever presses, (E) magazine entries and reward deliveries, and (F) ratios of magazine entries per active lever press and per reward. (G-J) Dopamine signals recorded during fibre photometry on (G-H) acquisition day 3, n=7 and (I-J) acquisition day 10, n=10. Photometry data presented shows average of all lever presses and magazine entries of all animals analysed. The grey area between −5 and −3 seconds pre-event was our baseline period, while the yellow area between 0 and 2 seconds post-event was the period in which we analyzed signals linked to reward behaviors.

### Fibre photometry of DA transients in the VP during instrumental reward learning

To characterize DA release in the VP during a self-administration paradigm in which animals learned to press an active lever to receive a sugar pellet delivered into the magazine, we performed fiber photometry recordings of DA transients using the GRAB-DA sensor ^40^. Four distinct phases of reward learning, including acquisition, extinction, reacquisition and progressive ratio test were recorded. Longitudinal recording of DA transients occurred from early and late (days 3 and 10) acquisition, early and late extinction (days 1 and 3), reacquisition test, and the motivation (progressive ratio) test. We obtained signals related to four behavioural actions: i) where an active lever press produced a sugar pellet reward within 0-2 sec of Lever press (LP+); ii) magazine entries on rewarded trials within 0-2 seconds of reward delivery (Mag+); iii) unrewarded active lever presses that occurred during time out period (LP−), where rats pressed the active lever during the 10-second time out period, and thus received no sugar pellet rewards; and iv) magazine entries during unrewarded trials (Mag−).

Fluorescence related to DA activity was quantified in two epochs: a two-second baseline period between 5 and 3 seconds before reward behaviours (−5 to −3 sec) and the two seconds immediately following a reward-related behaviour (lever press or magazine entry, 0 to 2 sec). The baseline period was established to minimise signals from preceding reward-related events. As there is a 10 second time out period between each LP+. There is no LP+ onset occurs, 10 seconds prior there is no LP+. As event-related dopamine responses were quantified within the first 2 s following behavioural onset, the −5 to −3 s baseline window provided a temporally isolated measure of ongoing fluorescence that was not influenced by preceding events-associated dopamine transients.

#### Acquisition

All rats successfully discriminated between the active and inactive levers from the first day of acquisition, and their instrumental responding became more robust across conditioning. This was confirmed by the statistical analysis, which found a significant linear increase in active lever presses across 10 days of acquisition training (F _(1,9)_ = 22.2, *p* = 0.001, see Figure 2E). By contrast, inactive lever presses remained consistently low and declined significantly across training days (F_(1,9)_ = 10.3, *p* = 0.011; mean (± SEM): day 1 = 13.3 (3.3), day 10 = 6.0 (1.7), see Figure 2E). The difference between active and inactive lever presses was also robustly significant (F_(1,9)_ = 47.1, *p* < 0.001, *d* = 1.0, 95% CI = 64.6, 128.0). Rats successfully discriminated between the active and inactive levers on day 3 (F_(1,9)_ = 6.3, *p* = 0.033, *d* = 1.1, 95% CI = 3.4, 64.2) and day 10 (F_(1,9)_ = 47.1, *p* < 0.001, *d* = 3.0, 95% CI = 64.6, 128.0), on the first of our recording days. However, the total number of active lever presses (F_(1,9)_ = 13.5, *p* = 0.005, *d* = 1.4, 95% CI = 23.8, 100.0), and the difference between active and inactive lever presses (F_(1,9)_ = 13.6, *p* = 0.005, *d* = 1.4, 95% CI = 24.1, 100.9) were both significantly less robust on day 3 than day 10, by which time rats had learned the reward rate and intertrial interval.

Fibre photometry data for day 3 and day 10 are shown in Figure 2G-J. Due to a technical malfunction, the data were lost for n = 3 rats, yielding a group size of n = 7 for acquisition day 3. On acquisition day 3, there were no significant event-related signals detected by the DA sensor upon specified behavioural onset compared to baseline (LP+: F_(1,6)_ = 2.3, *p* = 0.18; Mag+: F_(1,6)_ = 0.9, *p* = 0.38; LP−: F_(1,4)_ = 1.0, *p* = 0.38; Mag−: F_(1,6)_ = 0.9, *p* = 0.38). On acquisition day 10, we observed significant event-related DA signals compared to baseline. Significant DA spikes occurred after lever presses and magazine entries on rewarded trials (LP+: F_(1,9)_ = 39.6, *p* < 0.001, *d* = 2.9, 95% CI = 1.2, 2.5; ME+: F_(1,9)_ = 36.8, *p* < 0.001, *d* = 2.8, 95% CI = 1.1, 2.4).

By contrast, DA signals after lever presses on unrewarded trials were not significant (LP−: F_(1,9)_ = 0.1, *p* = 0.83), and signals after magazine entries on unrewarded trials were significantly negative (ME+: F_(1,9)_ = 10.3, *p* = 0.01, *d* = 1.2, 95% CI = −0.11, −0.64).

#### Extinction

Fibre photometry data during extinction behaviour are shown in Figure 3. Reduction of active lever pressing during extinction training was successful, as confirmed by a significant linear effect of day (F_(1,9)_ = 11.8, *p* = 0.007).

**Figure 3.**
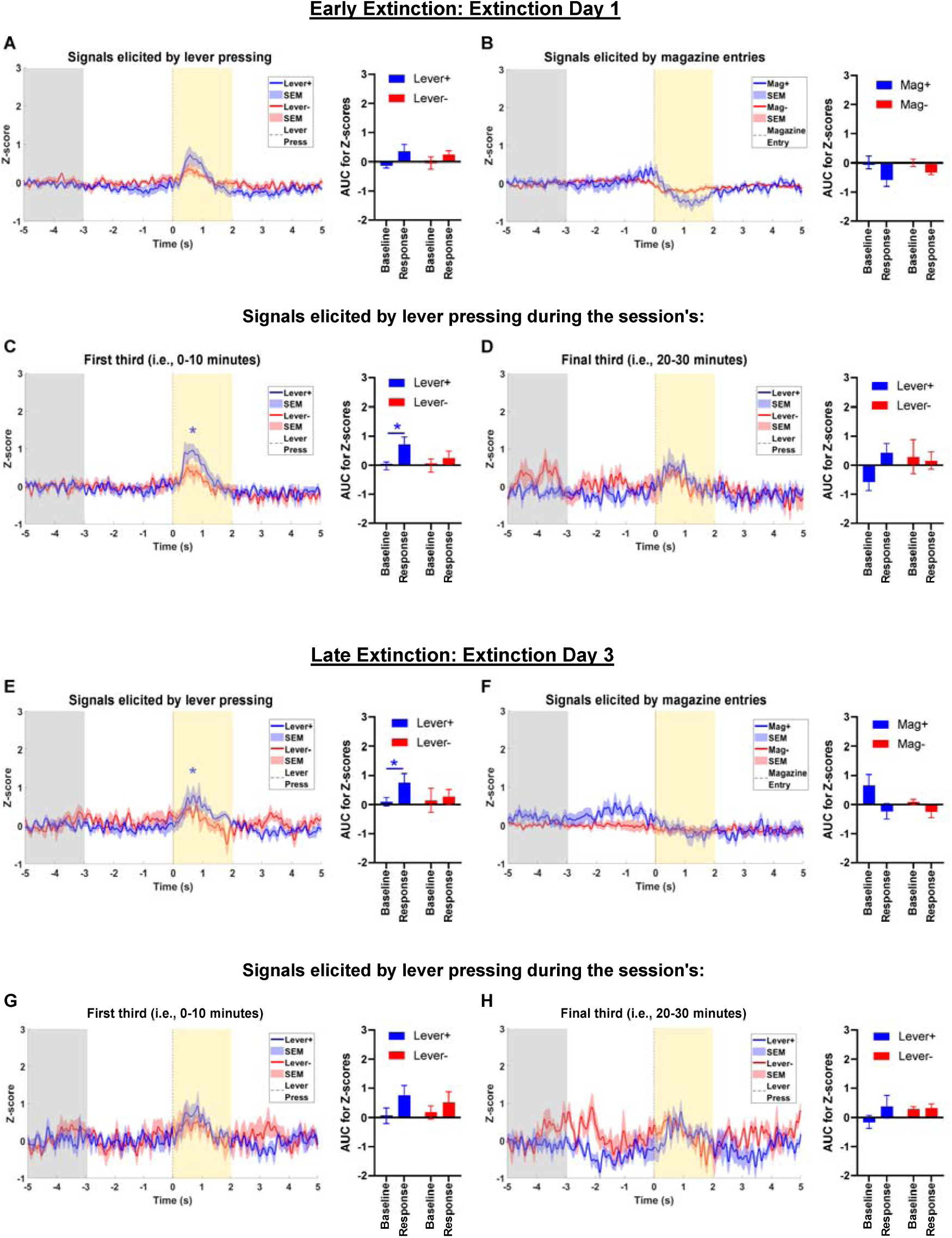
(A-B) Dopamine signals recorded by fibre photometry on extinction day 1. (C-D) Dopamine signals broken down into the (C) first 10 minutes and (D) third/final 10 minutes of extinction day 1. (E-F) Dopamine signals recorded by fibre photometry on extinction day 3. (G-H) Dopamine signals broken down into the (G) first 10 minutes and (H) third/final 10 minutes of extinction day 3. Photometry data presented shows average of all lever presses and magazine entries of n=10 animals. The grey area between −5 and −3 seconds pre-event was our baseline period, while the yellow area between 0 and 2 seconds post-event was the period in the grey area between −5 and −3 seconds pre-event was our baseline period, while the yellow area between 0 and 2 seconds post-event was the period in which we analyzed signals linked to reward behaviors.

Rats continued to discriminate between active and inactive levers on day 1 (F_(1,9)_ = 6.3, *p* =0.001, *d* = 2.1, 95% CI = 3.4, 64.2) and day 3 (F_(1,9)_ = 6.3, *p* = 0.06, *d* = 1.3, 95% CI = 3.4, 64.2). However, the overall number of active lever presses (F_(1,9)_ = 36.0, *p* < 0.001, *d* = 1.5, 95% CI = 34.0, 77.2), and the difference between active and inactive lever presses (F_(1,9)_ = 32.1, *p* < 0.001, *d* = 1.5, 95% CI = 28.8, 67.2), were significantly greater on day 1 than day 3. By contrast, magazine entries did not extinguish as the data exhibited no significant linear trend (F_(1,9)_ = 0.1, *p* = 0.76). These results demonstrate that instrumental responding extinguished, whereas magazine entries did not.

During extinction, rats continued to discriminate between previously associated lever with reward and magazine entry within 0 to 2 sec, (LP+ and Mag+) (LP− and Mag−). This allowed us to investigate how VP DA encodes the expectation, and surprising absence, of sugar rewards. Fibre photometry dopamine recording on extinction day 1, when averaging over the entire session, revealed no significant DA signals (LP+: F_(1,8)_ = 5.0, *p* = 0.06; Mag+: F_(1,8)_ = 4.1, *p* = 0.08; LP−: F_(1,8)_ = 1.0, *p* = 0.34; Mag−: F_(1,8)_ = 4.1, *p* = 0.08).

However, when analyzing the session in 10-min blocks, we found that LP+ triggered a significant DA spike in the first 10 minutes of the session (F_(1,8)_ = 7.0, *p* = 0.03, *d* = 1.2, 95% CI = 0.09, 1.38). As shown in Figure 3C-D, this LP+ DA signal was lost as the extinction session progressed, as it was absent in the third/final 10 minutes of the session (F_(1,7)_ = 3.5, *p* = 0.104). In contrast, signals for Mag+ and Mag−were nonsignificant in the first 10 minutes (Mag+: F_(1,8)_ = 1.9, *p* = 0.21; Mag−: F_(1,8)_ = 2.1, *p* = 0.18) and the third/final 10 minutes of the session (Mag+: F_(1,7)_ = 2.8, *p* = 0.16; Mag−: F_(1,6)_ = 0.5, *p* = 0.53). Photometry data from extinction day 1 are shown in Figure 3A-D.

On extinction day 3, DA signals were nonsignificant for Mag+ (F_(1,8)_ = 3.0, *p* = 0.12), LP− (F_(1,7)_ = 2.1, *p* = 0.66), and Mag− (F_(1,8)_ = 4.3, *p* = 0.07). Interestingly, the DA signal for LP+ was significant when averaging across the session (F_(1,8)_ = 6.3, *p* = 0.04, *d* = 0.9, 95% CI = 0.06, 1.25). However, when analyzing the first and third/final 10 minutes of the session, neither period contained significant DA signals (first 10 minutes: F_(1,7)_ = 2.6, *p* = 0.15; third/final 10 minutes: F_(1,6)_ = 2.8, *p* = 0.15), and the same was also true of the second 10 minutes (i.e., 10-20 mins; F_(1,7)_ = 3.0, *p* = 0.13). We suspect this is contributed by the low number of lever presses during later extinction. Photometry data from extinction day 3 are shown in Figure 3E-H.

#### Reacquisition

Behavioural data from reacquisition are shown in Figure 2D,E. Responses during reacquisition were significantly higher when compared to the final day of extinction; rats relearned to press the active lever at significantly higher rates (F_(1,9)_ = 12.1, *p* = 0.007), while pressing the inactive lever remained low and did not significantly increase (F_(1,9)_ = 3.3, *p* = 0.11). Photometry data from reacquisition are shown in Figure 4A-B. During reacquisition, lever presses and magazine entries on rewarded trials caused significant spikes in DA release (LP+: F_(1,9)_ = 27.5, *p* < 0.001, *d* = 2.9, 95% CI = 0.8, 3.1); Mag+: F_(1,9)_ = 10.9, *p* = 0.009, *d* = 1.5, 95% CI = 0.4, 1.9). On unrewarded trials, active lever presses triggered no significant DA activity (LP−: F_(1,9)_ = 1.0, *p* = 0.34), while magazine entries on unrewarded trials triggered a small, but significant, decline in DA (Mag−: F_(1,9)_ = 6.2, *p* = 0.04, *d* = 0.8, 95% CI = 0.02, 0.4). These data closely resemble the pattern we observed on acquisition day 10, as the signals were not significantly different between acquisition day 10 and reacquisition (LP+: F_(1,9)_ = 1.5, *p* = 0.252; LP−: F_(1,9)_ = 0.7, *p* = 0.418; Mag+: F_(1,9)_ = 3.2, *p* = 0.109; Mag−: F_(1,9)_ = 1.4, *p* = 0.270).

**Figure 4.**
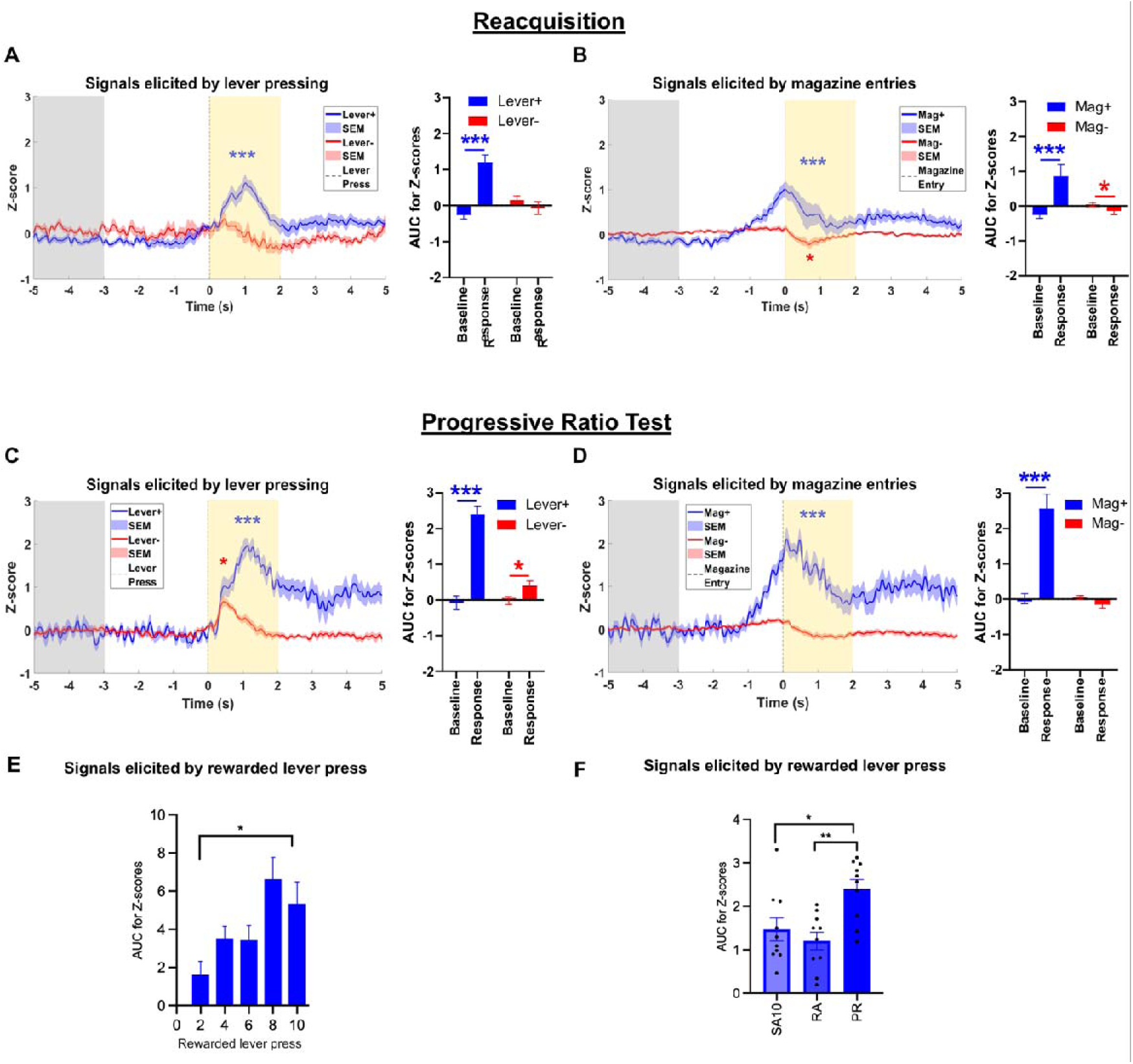
(A-B) Dopamine signals recorded by fibre photometry during reacquisition. (C-D) Dopamine signals recorded by fibre photometry during the progressive ratio test. Photometry data presented shows average of all lever presses and magazine entries of n=10 animals. The grey area between −5 and −3 seconds pre-event was our baseline period, while the yellow area between 0 and 2 seconds post-event was the period in The grey area between −5 and −3 seconds pre-event was our baseline period, while the yellow area between 0 and 2 seconds post-event was the period in which we analyzed signals linked to reward behaviors.(E) AUC of z-scores across reward lever presses during progressive ratio schedule (F) AUC of z-scores for LP+ during acquisition day 10, reacquisition and progressive ratio test.

#### Progressive ratio test

Behavioural data for the progressive ratio test are shown in Figure 2E. The ratio of rewarded lever presses increased linearly across the session. The average breakpoint was 25-33 active lever presses, beyond which rats gave up on lever pressing for sucrose pellets. During the progressive ratio test, this yielded an average of 8-10 sugar pellet rewards per rat.

Photometry data from the progressive ratio test are shown in Figure 4C-D. During the progressive ratio test, DA activity was significant during LP+ and Mag+ [(LP+: F_(1,9)_ = 54.4, *p* < 0.001, *d* = 3.2, 95% CI = 1.7, 3.2; Mag+: F_(1,9)_ = 38.8, *p* < 0.001, *d* = 2.6, 95% CI = 1.6, 3.5)]. Interestingly, on unrewarded trials, lever presses also caused a smaller, but still significant, spike in DA (LP−: F_(1,9)_ = 9.9, *p* = 0.01, *d* = 0.5, 95% CI = 0.1, 0.7), while magazine entries did not trigger significant DA activity (Mag−: F_(1,7)_ = 3.8, *p* = 0.08). Comparison between LP+ on the 1^st^ rewarded trial and the last LP+ shows significant increase in VP DA, F_(1,17)_ = 4.93, *p* = 0.040 (Figure 4E). Compared to acquisition day 10 and reacquisition, the signals observed on rewarded trials were significantly higher during the progressive ratio test (LP+: F_(1,9)_ = 86.4, *p* < 0.001, *d* = 1.5, 95% CI = 0.8, 1.3; Mag+: F_(1,9)_ = 15.6, *p* = 0.003, *d* = 1.3, 95% CI = 0.6, 2.2). By contrast, signals from unrewarded trials during the progressive ratio test were not different from acquisition day 10 and reacquisition (LP−: F_(1,9)_ = 2.4, *p* = 0.15; Mag−: F_(1,9)_ = 0.3, *p* = 0.61). One way ANOVA comparison of DA activity levels upon LP+ across acquisition day 10 (1.46 ± 0.26), reacquisition (1.20 ± 0.20) and progressive ratio test (2.39 ± 0.22), revealed significant differe42nce F_(2,18)_ = 7.39, *p* = 0.003. Bonferroni post hoc analysis reveals no group difference between acquisition and reacquisition day, *p =* 1.00. However significant difference between acquisition and PR task, *p =* 0.025 and reacquisition and PR task, *p* = 0.003 (Figure 4F)

### Histology

Bilateral cannula placements in the VP were confirmed using cresyl violet staining (see Figure 5A-B), yielding final groups of n = 8 for the vehicle group and n = 7 for the DA antagonist group.

**Figure 5.**
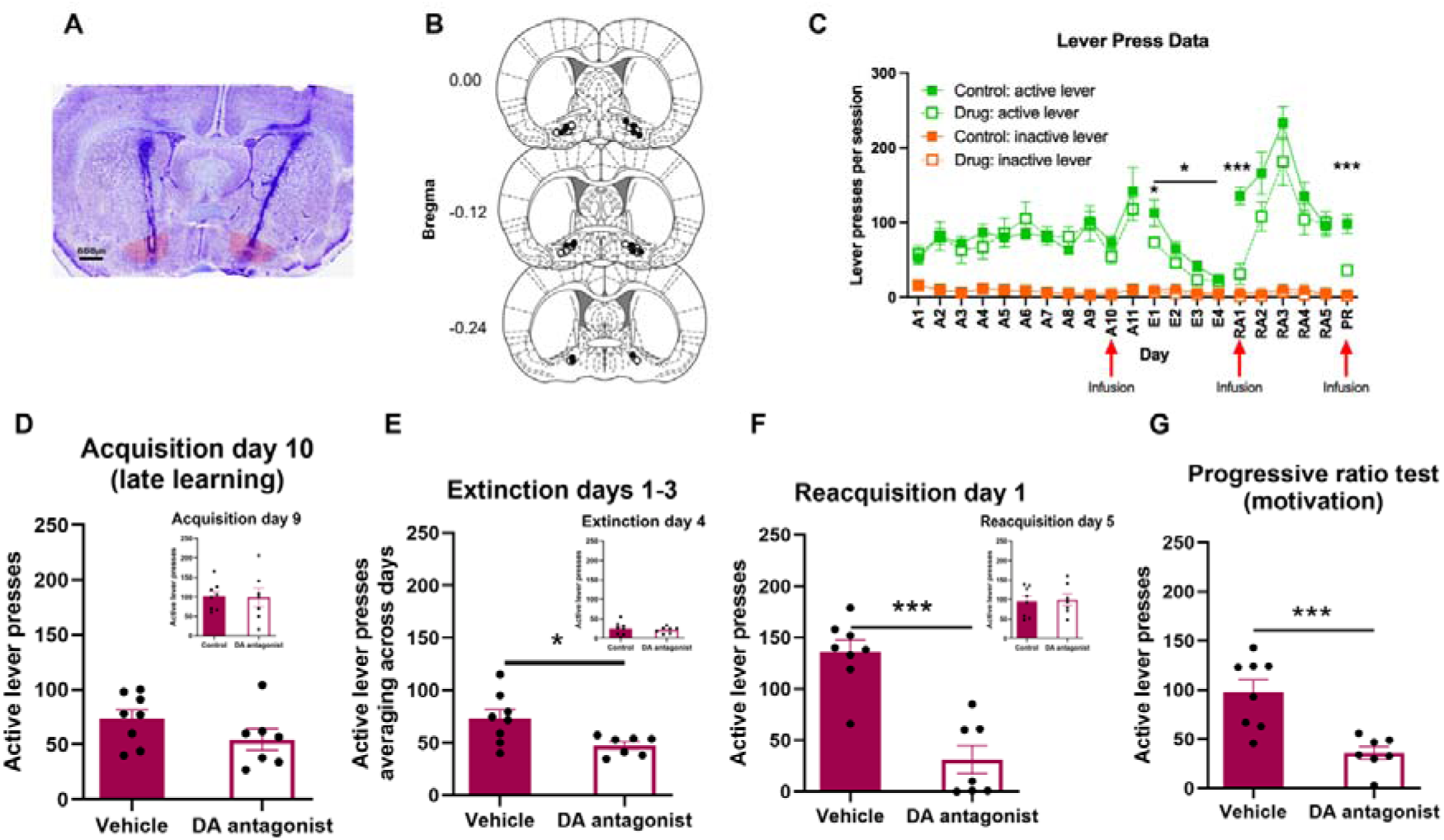
(A) Representative image of cannula placement in the VP and Histological reconstruction of cannula placements in the VP, n=7 vehicle, n=8 drug infusions.(B) Mean (± SEM) behavioral data for active and inactive lever presses during behavioral performance of pharmacological microinfusions into the VP. (C) Locomotor data showing distance travelled (D) Behavioral data of active lever presses on acquisition day 10 and an inset showing group matching on acquisition day 9. (E) Average active lever press during extinction days 1-3 and an inset showing group matching on extinction day 4. (F) Active lever presses during reacquisition day 1 and an inset showing group matching on reacquisition day 5. (G) Active lever presses during the progressive ratio test. All behavioral data are expressed as mean (± SEM).

### Behavioral pharmacology to confirm that VP DA activity is causal to reward behavior

The drug or vehicle was intracranially delivered to the VP on day 10 of acquisition, reacquisition, and the progressive ratio test. Notably, if a behavioural change was detected between groups upon vehicle or fluphenazine administration, animals were retrained to ensure that any effect of pharmacological manipulation did not impact subsequent behaviours.

### Behavior

#### Acquisition

The vehicle and DA antagonist groups successfully acquired instrumental responding to the active lever across conditioning, with no between-group differences from days 1 to 9. This was confirmed by the statistical analysis, which found a significant gradual increase of active lever presses from day 1 to day 9 of acquisition (main effect of day: F_(1,13)_ = 19.7, *p* = 0.001; main effect of group: F_(1,13)_ = 0.02, *p* = 0.89), while inactive lever presses remained low and declined further across training days (main effect of day: F_(1,13)_ = 28.3, *p* < 0.001; main effect of group: F_(1,13)_ = 0.7, *p* = 0.43). Consequently, the difference between active and inactive lever presses was robustly significant on acquisition day 9 (main effect of lever type: F_(1,13)_ = 56.3, *p* < 0.001, *d* = 2.8, 95% CI = 68.1, 123.2; main effect of group: F_(1,13)_ = 0.03, *p* = 0.87; lever type x group interaction: F_(1,13)_ = 0.003, *p* = 0.96).

Immediately before the acquisition session on day 10, rats received intra-VP infusions of the vehicle or the DA receptor antagonist, fluphenazine. The DA antagonist did not acutely suppress reward behaviours (see Figure 5D), as there were no between-group differences for active lever presses (F_(1,13)_ = 2.2, *p* = 0.16) or magazine entries (F_(1,13)_ = 0.8, *p* = 0.40).

#### Extinction

Active lever pressing was successfully extinguished during extinction training, as confirmed by a significant linear effect of day (active lever presses: F_(1,13)_ = 86.7, *p* < 0.001). These data also revealed a drug effect from acquisition day 10, as the DA antagonist group pressed the active lever significantly less than the vehicle group, when averaging across the first three days of extinction (F_(1,13)_ = 5.1, *p* = 0.04, *d* = 0.91, 95% CI = 1.0, 39.1, see Figure 5E). Group comparison on extinction day 1 t_13_ = 1.96, *p*= 0.036. Repeated measure within animal comparison over acquisition day 10, 11 and extinction day 1 show no significant difference between performance during acquisition and extinction; F_(2,28)_ = 2.31, *p* = 0.24, F_(2,28)_ = 2.86 *p* = 0.11.

Magazine entries also partially extinguished, but with no sign of a group difference (effect of trials: F_(1,13)_ = 38.1, *p* < 0.001; effect of group: F_(1,13)_ = 3.0, *p* = 0.11). This suggests that blocking DA activity in the VP on acquisition day 10 disrupted the learned association between active lever presses and sugar rewards. By extinction day 4, however, the groups were no longer significantly different (see Figure 5E), as confirmed by the absence of between-group differences in active lever pressing (F_(1,13)_ = 0.2, *p* = 0.67) and a significant day x group interaction for active lever presses (F_(1,13)_ = 4.9, *p* = 0.045, 95% CI = 0.7, 54.5).

#### Reacquisition

Following extinction, rats underwent reacquisition, with infusions of vehicle or fluphenazine immediately before the session. During reacquisition, the DA antagonist group made significantly fewer active lever presses (F_(1,13)_ = 33.8, *p* < 0.001, *d* = 3.0, 95% CI = 65.7, 143.3, see Figure 5F), inactive lever presses (F_(1,13)_ = 10.3, *p* = 0.007, *d* = 1.7, 95% CI = 1.5, 7.9), and magazine entries (F_(1,13)_ = 8.7, *p* = 0.011, *d* = 1.5, 95% CI = 1.5, 7.9) than the vehicle group. Before moving on to the progressive ratio test, rats were then retrained until both groups were matched for reward learning. Group comparison of the reacquisition day lever presses after infusion on reacquisition test did not reveal any group difference; t_13_ = 1.650, *p* = 0.061. Additional sessions of reacquisition training (see Figure 5F), at which point there were no between group differences for active lever presses (F_(1,13)_ = 0.2, *p* = 0.84), inactive lever presses (F_(1,13)_ = 0.02, *p* = 0.88) or magazine entries (F_(1,13)_ = 1.7, *p* = 0.21) to ensure no group difference in performance during progressive ratio test.

#### Progressive ratio test

Rats were infused with vehicle or fluphenazine immediately before the progressive ratio test. Antagonizing DA receptors in the VP significantly reduced lever press for sugar rewards, as evidenced by a main effect of group for active lever presses (F_(1,13)_ = 17.3, *p* = 0.001, *d* = 2.2, 95% CI = 29.7, 94.0, see Figure 5G), and a significantly lower breakpoint for DA antagonist group (F_(1,13)_ = 12.2, *p* = 0.004, *d* = 1.8, 95% CI = 4.0, 17.1). While latency to first lever press remain unaffected, t_13_= 0.43, *p* = 0.337, reduction in motivation was evident by significant decrease magazine entries t_13_= 2.12, *p* = 0.02.

### Locomotor

Locomotor analysis of VP was conducted in a separate cohort of rats and only assessed on locomotor behaviour n=10. Animals habituated one day before micro-infusions. Animals were infused in a counterbalanced manner across 2 days with vehicle and drug infusions. Data presented at cm (mean ± SEM), vehicle = 28246.32 ±8345.20 and drug group = 44189.07 ±50035.93. Paired sample tests analysis t_9_ =- 0.633, *p* = 0.262. Locomotor activity is presented as distance travelled in 30mins.

## Discussion

The VP has been established as an important node for reward processing, reward prediction, and motivation^19,25,28,41,42^, here we demonstrate that DA contributes to these processes. We found significant DA activity in the VP during established learning, extinction, reacquisition, and the progressive ratio test. Our results show that the role of VP DA is multifaceted, as it supports instrumental reward learning and drives motivation to work for rewards.

The DA mesolimbic pathway has been widely studied, with most examining DA projections to the Acb in reward processing. A number of the accumbal DA afferents send collaterals to VP^35^. We also show a separate population of DA neurons that project only to the VP (Figure 1). Other studies have found DA modulation in VP often has more intense effects than in the accumbens^43,44^. A series of papers using neural recording with electrode arrays in the Acb and VP reported a significant proportion of VP neurons respond earlier than Acb to rewards^26^.

While roles of afferent inputs from the Acb, subthalamic nucleus and lateral hypothalamus to the VP well established^16,17,41,45–47^, considerably less attention has been given to DA efferents. Our findings build on these fundamental studies, showing that reward-related VP activity depends crucially on local DA release during acquisition, reacquisition, and motivated behaviors.

### *VP DA* supports Model-Based Reward Learning and Action-Outcome Associations

Our findings indicate that DA release within the VP contributes to the formation and updating of action-outcome associations during instrumental reward learning. While there was no significant DA release during early acquisition, by late acquisition, the VP DA signal was robustly and significantly increased following rewarded lever presses and magazine entries, and the same pattern was subsequently observed during reacquisition. These findings imply that VP DA activity scales with the strength of the action-outcome association. The strengthening of VP dopamine responses observed across early and late acquisition therefore likely reflects increasing confidence in the action-outcome relationship. Furthermore, blockade of dopamine receptors within the VP impaired subsequent reacquisition and reduced responding during early extinction, suggesting that VP contributes to the maintenance and updating of learned reward contingencies.

Importantly, these observations are parallel to findings from other studies showing that VP neurons encode the association between reward-seeking behaviour rather than reward delivery alone^42^. Together with our pharmacological findings, these data suggest that DA release within the VP may provide a teaching signal that reinforces and updates associative representations encoded by VP neurons.

This interpretation that VP neurons encode reward association is consistent with evidence that DA signals support model-based representations of reward contingencies alongside classical reinforcement learning mechanisms^1,48^. Recent work has demonstrated that mesolimbic dopamine activity reflects inferred relationships between cues, actions, and outcomes, allowing animals to adapt behaviour when contingencies change rather than merely reinforcing previously rewarded actions ^3,4^.

### VP DA Tracks Expectation of Reward

Significant VP DA activity occurred at the beginning of extinction day 1 session but insignificant across the session. Early in the first extinction session, DA signals for previously rewarded lever (Lever+) were significant. However, not during the Lever-signal showed that the rats successfully discriminated between trial types without reward deliveries. This suggests that the significant Lever+ signal during extinction day 1 likely reflects rats’ shifting expectation of reward delivery across the extinction session. These results from extinction day 3 are less clear but suggestive of similar conclusion. The is also detected by the lack of significant DA activity when rats no longer expected reward delivery during the end of the extinction session. Hence, our extinction data suggest that VP DA reflected the expectation of reward.

#### VP Dopamine Encodes Reward Prediction and Prediction Error Signals

A central question arising from the present findings is whether DA release within the VP contributes to reward prediction error (RPE) signalling. Although single-unit recordings in the VP have demonstrated that a subset of VP neurons respond to RPE signals, where VP RPE signals emerged rapidly and were capable of encoding both positive and negative prediction errors^25^.

Here several aspects of our data support that VP DA participates in reward prediction processes. During late acquisition and reacquisition, DA transients occurred following rewarded lever presses and magazine entries, consistent with positive reward prediction signals. Conversely, during extinction, DA responses were strongest early in the session when animals still expected reward delivery and diminished as extinction progressed and reward expectations were updated. This temporal shift in DA signals upon changes in reward expectations is consistent with reward prediction process.

Interestingly, we observed the clearest evidence of prediction related signalling during the progressive ratio task. Under this schedule, the relationship between action and reward becomes increasingly uncertain because reward delivery is progressively delayed by escalating response requirements. The emergence of DA transients following expected but unrewarded lever presses suggests that VP DA may encode an expectation of future reward despite the immediate absence of reinforcement.

Rather than reflecting a classical negative prediction error, these signals may represent a prospective reward expectation signal that maintains behavioural engagement under conditions of increasing effort. As seen within the progressive ratio schedule, comparison between the 1^st^ and last rewarded lever press were significantly different (Figure 4E). Such a mechanism would be consistent with theoretical frameworks proposing that DA contributes not only to reward error but also to during uncertain or changing contingencies.

#### VP DA is Augmented by Motivation

Modern theories increasingly distinguish DA role in learning and motivation^6^. Others have consistently demonstrated DA is essential for effort-based decision-making and willingness to work for rewards^49^. The DA signal was very similar on acquisition day 10 and reacquisition but significantly higher during the progressive ratio test. Therefore, the DA enhancement detected between the first and last rewarded response during progressive ratio task therefore cannot be solely explained by reward outcomes and instead suggests a role for VP dopamine in motivational processes. During progressive ratio performance, DA activity increased as response requirements progressively escalated and significantly elevated relative to standard fixed-ratio schedules. Moreover, VP DA receptor blockade significantly reduced active responding during progressive ratio task. These observations suggest that VP DA may increase the incentive value of rewards when greater effort is required to obtain them. The present findings extend these concepts of DA functions in the VP.

These findings also align closely with the work of Ottenheimer and colleagues demonstrating that VP neurons encode incentive salience and relative reward value. In their studies, VP neuronal activity tracked the motivational significance of reward-predictive cues and often encoded reward value earlier and more robustly than nucleus accumbens neurons. The heightened DA release observed during progressive ratio task may therefore represent a neuromodulatory mechanism through which motivational value signals are amplified within the VP. Together, these findings support an emerging model in which the VP functions as a critical hub integrating reward prediction and motivational state.

### Limitations of the study

Several limitations should be considered when interpreting the present findings. First, although fluphenazine provided a useful approach for establishing a causal role of VP DA signalling, it is a non-selective dopamine receptor antagonist. The application of a non-specific DA antagonist was based on recent transcriptional analysis in rodents and human VP neurons that shows expression of DA receptors 1-3^9,23^. The dose of fluphenazine selected was based on application of effective dose in previous work using self-administration paradigm^38^. Although the concentration of Fluphenazine was quite low, there may be confounding effects on 5-HT2A and Histamine receptors ^38,50^.

Second, although our findings are consistent with RPE related processes, the behavioural design was not specifically optimized to dissociate reward prediction, reward expectancy. Hence interpretation of the progressive ratio data would be better assessed using classical reward prediction tasks. The elevated DA signals observed during progressive ratio task on unrewarded lever presses may reflect reward prediction errors, persistent reward expectation, increased motivational drive, or a combination of these processes. Dedicated tasks involving probabilistic reward delivery, outcome devaluation, or reward omission, would allow more precise identification VP dopamine release functions

Finally, while VP has been shown to play a central role in the reinforcing effects across a variety of drugs, in this paradigm sugar pellets were used as reinforcement^30^. Given that both natural rewards and substances of abuse engage overlapping mesolimbic DA circuits, it is plausible that VP DA signalling contributes to reward processing across multiple reward modalities. Future studies should therefore examine VP DA release during self-administration of drugs of abuse.

## Conclusions

Despite these limitations, the convergence of anatomical tracing, longitudinal dopamine recordings, and causal pharmacological manipulations provides strong evidence that DA signalling within the VP contributes to reward learning, reward expectation, and motivational processes. These findings establish VP DA as an important component of the broader mesolimbic reward network.

## Author contribution

A.A.P. and JLC conceptualised the project, interpreted the data for the work, and edited the manuscript. A.A.P. and M.W.S. conducted the experiments and prepared the manuscript.

## Acknowledgement

This work was supported by the Australian Research Council (DP220100412) to A.A.P and J.L.C.

The authors declare no competing financial interests.

